# Acoustic target strengths and swimbladder morphology of Chub mackerel *Scomber japonicus* in the Northwest Pacific Ocean

**DOI:** 10.1101/2024.05.03.592349

**Authors:** Euna Yoon, Hyungbeen Lee, Yong-Deuk Lee, Yong Jin Choo, Jeong-Hoon Lee, Jung Nyun Kim

## Abstract

The Northwest Pacific chub mackerel (*Scomber japonicus*) is one of the most productive, economically important fishery resources worldwide. Due to fluctuations in their abundance and distribution, there is a pressing need to accurately assess this species and to ensure total allowable catch limits are followed. Acoustic target strength (TS; dB) measurements of *Scomber japonicus* were conducted at 38, 70, and 120 kHz using a split-beam echosounder of individuals from nine size groups (mean fork length, 10.8∼28.3 cm) swimming freely in a net cage within a seawater tank. An underwater camera was utilized to simultaneously measure the swimming angle. A least-squares regression analysis revealed that when the slope was constrained to 20, as per the generally applicable morphometric equation, the resulting values for the constant term (*b*_*20*_) were ‒67.7, ‒66.6, and – 67.3 dB at 38, 70, and 120 kHz, respectively. The mean swimming angle of *S. japonicus* across the groups was –10.5∼9.6° (standard deviation (SD), 16.3∼33.3°). In addition, the ratio of swimbladder height to swimbladder length, swimbladder length to fork length, and tilt angle of the swimbladder (mean ± SD) were 0.191 ± 0.060, 0.245 ± 0.055, and 9.6 ± 3.0°, respectively. These results can be used for the acoustic stock assessment of *S. japonicus* in the Northwest Pacific Ocean.

## 1. Introduction

The chub mackerel (*Scomber japonicus*), belonging to the mackerel family (*Scombridae*), is widely distributed in temperate marine areas including Korea, Japan, the East China Sea, and the eastern Pacific Ocean [1–4]. It is one of the most productive and economically important fisheries resources in the world, with annual catches reaching ca. 1.3 million metric tons [5]. The abundance of the Pacific stock of chub mackerel declined in the 1990s and early 2000s. Since 2013, when a strong year class occurred, its abundance dramatically increased [6,7]. In Korea, chub mackerel has consistently recorded the second-highest catch volume after anchovy, with an average annual catch of 110,640 metric tons from 2018 to 2022 [8]. Consequently, total allowable catch (TAC) limits were implemented for the sustainable management of chub mackerel in Korea, Japan, and China [9–12].

Research on pelagic fish using acoustic techniques involves collecting data from scientific echo sounders installed on ships proceeding along planned acoustic transects in the survey area, while simultaneously conducting catch surveys to confirm the species identification and length distribution data derived from the echo sounders. This information is then used to estimate the distribution and abundance of the target species [13]. For the identify and quantify of target species in the survey area, the average acoustic target strength (TS, dB) of the target species is essential [14]. This parameter is generally expressed as TS = 10·log_10_σ / 4π [15], where σ is the backscattering cross-section of the fish. TS length (L, cm) can be expressed as TS = *a* log_10_(*L*) + b, where the slope *a* and the intercept *b* (dB) are generally assumed to be species-specific constants. When the backscattering cross-section is proportional to the length squared, *a* is normally close to 20; thus, TS = 20·log_10_(*L*) + *b*_20_, where *b*_20_ is the estimated intercept given a slope of 20.

Pelagic fish TS data are obtained either through *ex situ* experiments in a seawater tank or *in situ* at sea, in conjunction with catch surveys, and are validated by comparison with a theoretical model to derive the TS-L relationship for the target species [16–18]. Most *ex situ* experiments on chub mackerel TS have been conducted using dead specimens [19,20]; however, it may result in inflated values compared to living specimens due to the expansion of body tissues or presence of gas bubbles [21]. Moreover, previous *ex situ* experiments have been limited to specimens with a total length of − 23 cm [19]. Recently, chub mackerel TS has also been studied using the theoretical Kirchhoff-ray mode (KRM) model [7,22], but deriving an accurate and reliable TS-L relationship requires experiments to be conducted on living specimens of various sizes in a free swimming state.

Factors affecting fish TS signals underwater include swimming posture angle, detector operating frequency, distribution depth, sound speed ratio, density ratio, and the presence and morphology of the swimbladder. In particular, the TS signal of fish with a swimbladder depends on the large density ratio difference between gas and water by more than 90% [23-25]. Therefore, when studying the TS-L relationship in Pacific chub mackerel, the morphology of its swimbladder must be taken into account during the analysis and interpretation of TS signals. In this study, we measured the *ex situ* TS at 38, 70, and 120 kHz for various length distribution groups of living chub mackerel and simultaneously filmed the swimming posture angle to verify fish movement. Subsequently, we investigated the characteristics of the swimbladder to validate the TS signal results.

## 2. Material and methods

### *Ex situ* target strength measurement

The chub mackerel TS experiments were conducted from April to September 2022 at the Fisheries Resources Research Center’s seawater acoustic tank (5 m length × 5 m width × 10 m depth) (Fig 1) at the National Institute of Fisheries Science. Specimens used in the experiment were caught during a set-net survey off the coast of Tongyeong, Gyeongsangnam-do (34° 47.0′N, 128° 26.4′E) and transported to the laboratory in a seawater tank, where they were acclimated in a circular tank (2.75 m diameter × 0.8 m depth) for 24 h prior to the TS experiment.

**Fig. 1.**
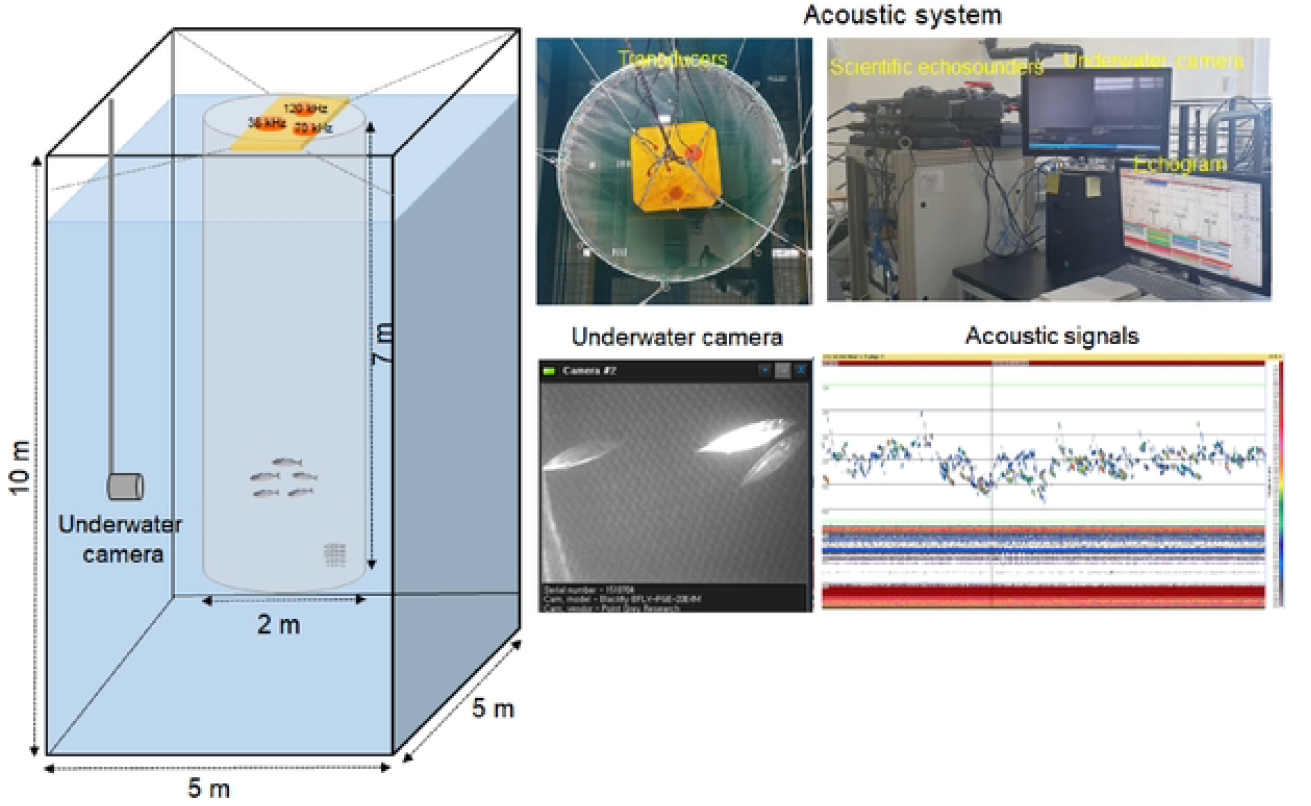
Experimental setup for the target strength measurements of chub mackerel (*Scomber japonicus*) using a scientific echosounder at 38, 70, and 120 kHz.

The system consisted of a SIMRAD EK80 split-beam scientific echosounder (SIMRAD, Kongsberg Maritime AS, Kongsberg, Norway). It operated at frequencies of 38 kHz (ES38-10), 70 kHz (ES70-7C), and 120 kHz (ES120-7C), with a transducer utilizing –3 dB beam widths of 10°, 7.1°, and 7.1°, and respective power outputs of 1500 W, 750 W, and 250 W. The pulse duration and ping rate for all frequencies were set to 0.256 ms and 2 pings/s, respectively. Before the experiment, the temperature and the salinity of the seawater in the acoustic tank were measured with an YSI 30M instrument (YSI, Yellow Springs, OH, USA) to calculate sound speed [26]. Additionally, the transducers were calibrated with a 38.1-mm tungsten carbonate standard sphere [27].

For the TS experiment, a cylindrical cage (2 m diameter × 7 m depth, equipped with a stainless steel ring and 1.0-cm mesh nylon netting) was installed in the tank to streamline sample replacement and to accommodate the beam width of transducers for each frequency (Fig 1). Transducers were installed in the tank vertically at a depth of 30 cm. For the experiment, visually similar sizes of chub mackerel specimens acclimated in the circular tank were collected in small buckets without exposing individuals to air and transported to the cage installed in the acoustic tank, where the TS signals of freely swimming specimens were acquired. During the experiment, an underwater camera was installed on the side of the tank to observe swimming posture angle. The camera was vertically adjustable and recorded as it was positioned to the depth layer where the fish were primarily distributed. After the TS experiment, the fork length (FL) and wet weight (W) of the specimens were measured.

Chub mackerel specimens were placed in the cage divided into nine groups of similar length, and an effort was made to visually ensure that lengths between groups did not overlap. The average FL for the groups ranged from 10.8<u>–</u>28.3 cm, with a length range within groups of 0.2–1.1 cm (Table 1).

**Table 1.**
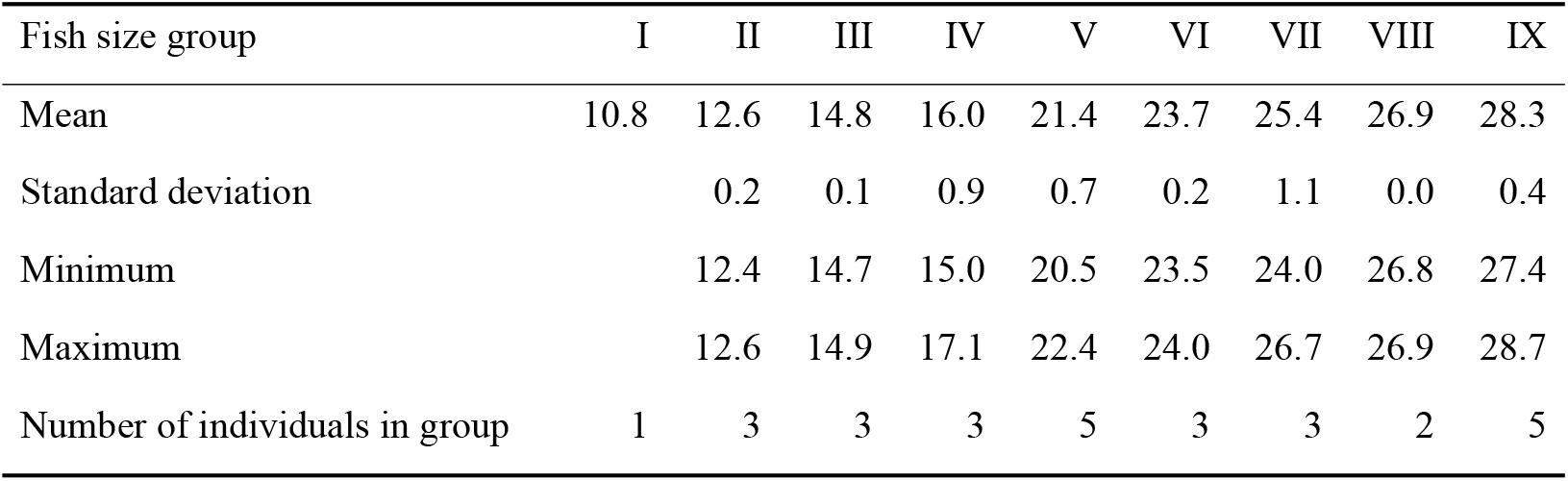
Characteristics of size groups of chub mackerel used in the cage experiments (fork length in cm).

### Acoustic data analysis

Group-specific TS acoustic signal analysis was conducted using the Echoview Software, version 11.0. TS signal analysis involved the extraction of individual fish signals using single echo detection (SED) and detection of fish track modules after removing basic noise like bottom reflection (Fig 1). The single echo detector parameters were set to accept echoes with a threshold value of −65 dB. The range for SED was set to minimum and maximum pulse lengths 0.7 and 1.5, respectively, a maximum beam compensation of 6 dB, and a maximum standard deviation for minor and major axis angles of 0.6°. For detecting fish tracks, the minimum number of single targets in a track was set to 3, the minimum number of pings in a track was set to 3, and the maximum gap between single targets was set to 1 ping. The average TS values for each size group of chub mackerel were calculated using the SED and detect fish tracks modules, with mean TS = 10·log_10_ (Σσ / *n*_i_), where σ is the backscattering cross-section of the fish and *n* is number of specimens.

### Swimming angle measurement

For the analysis of swimming posture angles, video data from underwater cameras were converted into images at 1-second intervals, and the swimming posture angles were measured using ImageJ software (National Institute of Health, USA) based on the center of the fish’s snout and tail fin. Swimming posture angles were defined as positive angles (+) when the fish’s head was pointing upwards and as negative angles (<u>−</u>–) when pointing downwards. Among the video data of the nine groups, group 1 (a single specimen) was not recorded due to a camera depth setting error, so swimming posture angles were analyzed for the remaining eight groups only.

### Swimbladder characteristics

To investigate the swimbladder morphology of the chub mackerel used in the TS experiment, samples were subjected to X-ray imaging. After the TS experiment, samples measured for fork length (L, cm) and wet weight (W, g), and then rapidly frozen in a freezer at below −40°C−after placing them in bottles with seawater. Frozen samples for X-ray imaging were slowly thawed in cold water for 24 hours in their bottles to minimize deformation of the swimbladder’s shape [28]. The lateral and dorsal aspects of all individuals used in the TS experiment were photographed using soft X-ray (150 kV, 2mA; SOFTEX M-150W, SOFTECS Corporation, Tokyo, Japan) (Fig 2). Among the 28 chub specimens used in the TS experiment, the swimbladder of one individual (FL = 12.5 cm) from group 2 was damaged and was omitted from analysis. Swimbladder morphology was characterized by tilt angle (SBA), length (SBL), height (SBH), and width (SBW) (Fig 2), and the equivalent radius (α_esr_ = (a × b × c)^1/3^) was calculated using the semi-major axis (a = SBL/2) from the lateral aspect, the semi-minor axis (b = SBH/2), and the dorsal aspect (c = SBW/2) [25].

**Fig 2.**
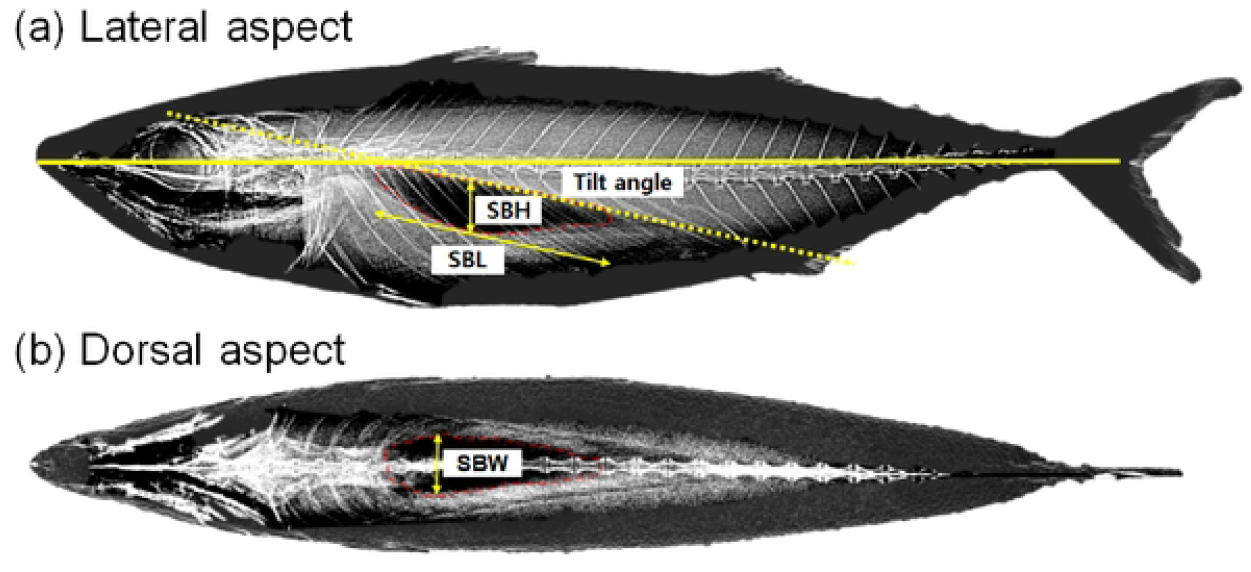
Soft X-ray images of the (a) lateral and (b) dorsal aspects of *S. japonicus*. The red lines are the boundary of the swimbladder. The swimbladder angle (θ) indicates the tilt angle of the swimbladder relative to the centerline between the anterior and posterior margins. SBH, SBL, and SBW refer to swimbladder height, swimbladder length, and swimbladder width, respectively.

## 3. Results

### Environment and specimens

During the TS experiments, the seawater tank’s temperature and salinity ranged from 16.1<u>–</u> to 23.8°C−and 29.2<u>–</u>33.4 psu, respectively, resulting in sound speeds ranging from 1508.3–1525.1 m/s. A regression model fit to the relationship between FL and W for the 28 chub mackerel used in the experiment was W = 0.0029·FL^3.4537^ (R^2^=0.99) (Fig 3), and mean FL showed significant between-group differences (ANOVA, p<0.05).

**Fig 3.**
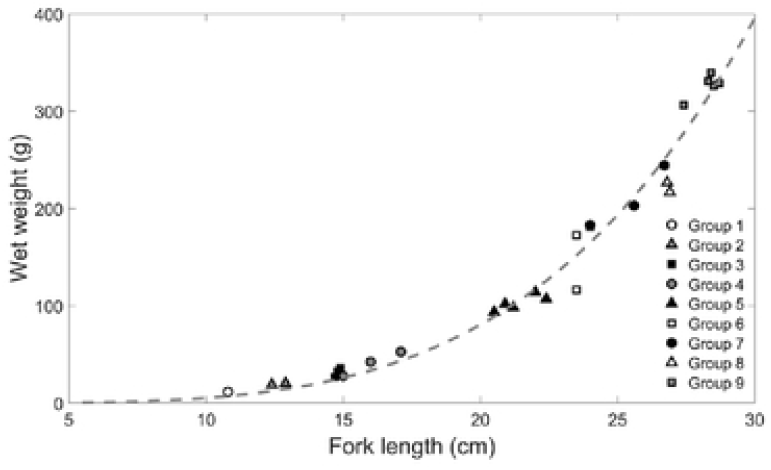
The relationship between fork length (FL, cm) and wet weight (W, g) of nine groups of *S. japonicus*.

### *Ex situ* target strength and swimming angle

The range of TS for the 9 groups of chub mackerel was −63.5 to −28.7 dB at 38 kHz, −64.2 to −27.7 dB at 70 kHz, and −64.1 to −29.2 dB at 120 kHz. TS per group mostly followed a Gaussian distribution (Fig 4). The variation in TS per group was 9.2–32.3 dB at 38 kHz, 9.9–30.9 dB at 70 kHz, and 18.9–33.7 dB at 120 kHz. The range of TS variation per group at 38 kHz and 70 kHz increased with the length of FL (R^2^ = 0.56∼0.94, p>0.05), while at 120 kHz, there was no correlation between size by FL and TS (R^2^ = 0.50, p<0.05). The mean TS per group was −45.8 to −35.8 dB at 38 kHz, −46.6 to −35.9 dB at 70 kHz, and −47.5 to −36.6 dB at 120 kHz, with all frequencies displaying concurrent increases in TS and body size (paired *t*-test, p<0.05). Notably, the correlation coefficients of TS by size per frequency showed no significant differences (paired *t*-test, p>0.05) (Fig 5).

**Fig. 4.**
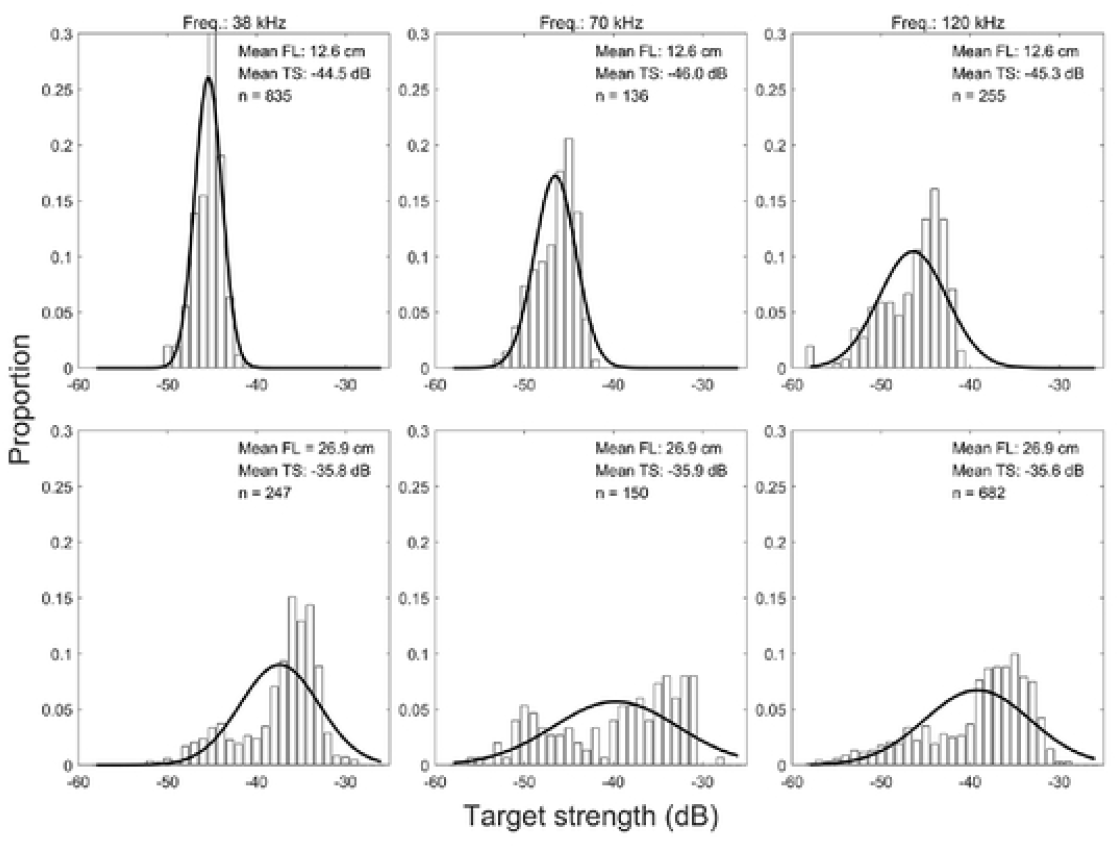
Target strength (dB) distribution of the mean of fish tacks based on three transducers (indicated on top) and in fish size groups 2 (mean FL: 12.6 cm) and 8 (mean FL: 26.9 cm) (indicated in panels). The black line shows the estimated probability density function (PDF).

**Fig. 5.**
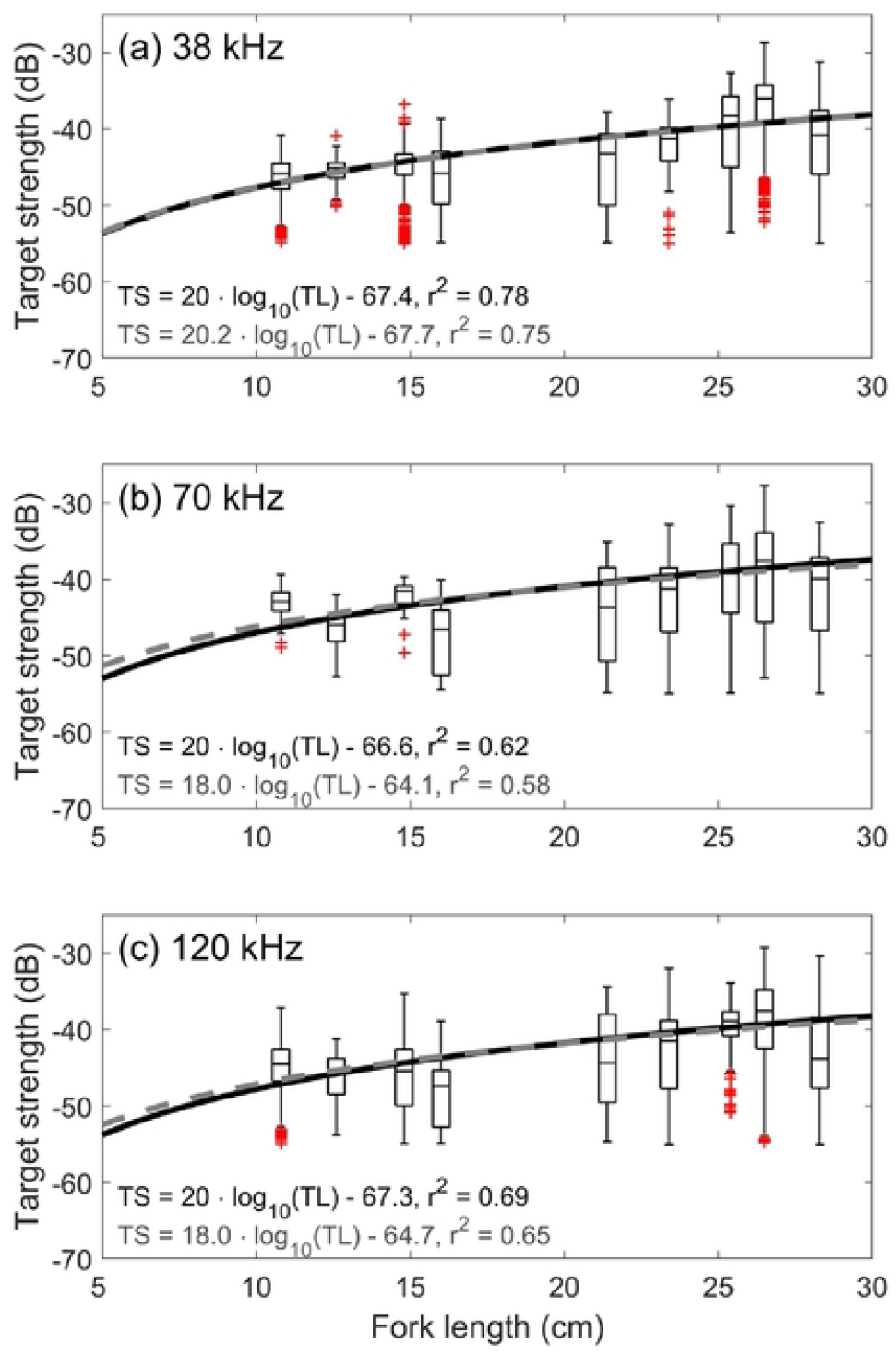
Relationship between the target strength (TS) and mean fork length of fish groups (cm) at (a) 38, (b) 70, and (c) 120 kHz. The results of the standard linear regression model (gray dot line) and those obtained with the slope forced to 20 (black dashed line) are also reported.

Least-square regression models of TS vs log(FL) by acoustic frequency across all groups were fit as follows:

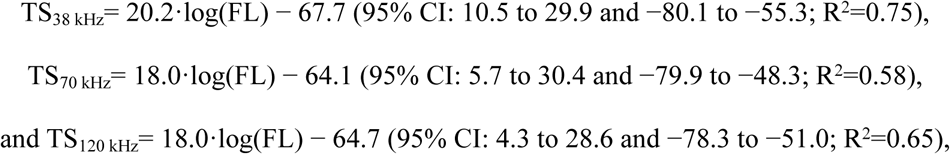

giving the confidence intervals for *a* and *b*, respectively.

When forcing the model fit to a slope of 20, as per the standard formula, the following models were estimated:

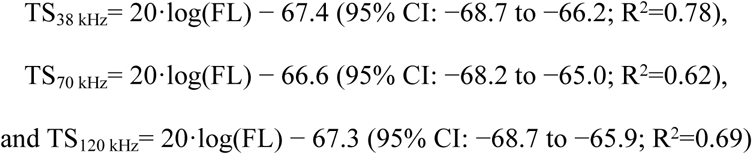

The group-specific TS of chub mackerel at 38, 70, and 120 kHz frequencies at *a* = 20 were similar, with b_20_ within a range of 0.9 dB (highest for 70 kHz and lowest for 120 kHz).

The swimming posture angles of chub mackerel groups showed a wide range of angles ranging from −71.4–69.2°, with group means of −10.4–9.6° and standard deviations of 16.3–33.3° (Fig 6). The distribution of swimming posture angles was mostly unimodal, with the larger-sized groups 8 and 9 displaying bimodal distributions. The correlation between mean body length and swimming posture angle was negative and differed significantly between groups (R^2^=0.61, paired *t*-test, p<0.05).

**Fig 6.** Swimming angle distributions of chub mackerel in size groups 2∼9 obtained from lateral view by underwater camera during the TS experiment. No group 1 data were captured due to the small number of fish (single specimen).

### Swimbladder morphology

The length of the chub mackerel swimbladders as imaged by X-ray ranged from 1.6–8.6 cm (mean ± SD: 5.5 ± 2.3 cm), the height from 0.2–2.3 cm (1.1 ± 0.6 cm), and the width from 0.1–2.4 cm (1.1 ± 0.5 cm), all showing a proportional increase with size (paired *t*-test, p < 0.01) (Fig 7). The equivalent radius of the swimbladders ranged from 0.20–1.76 cm (0.92 ± 0.42 cm), showing a positive correlation with length. Contrastingly, the tilt angle of the swimbladders ranged from 6.0–19.0° (9.6 ± 3.0°; n = 27), displaying a negative correlation with body length (paired *t*-test, p<0.01, R^2^=0.06).

**Fig. 7.**
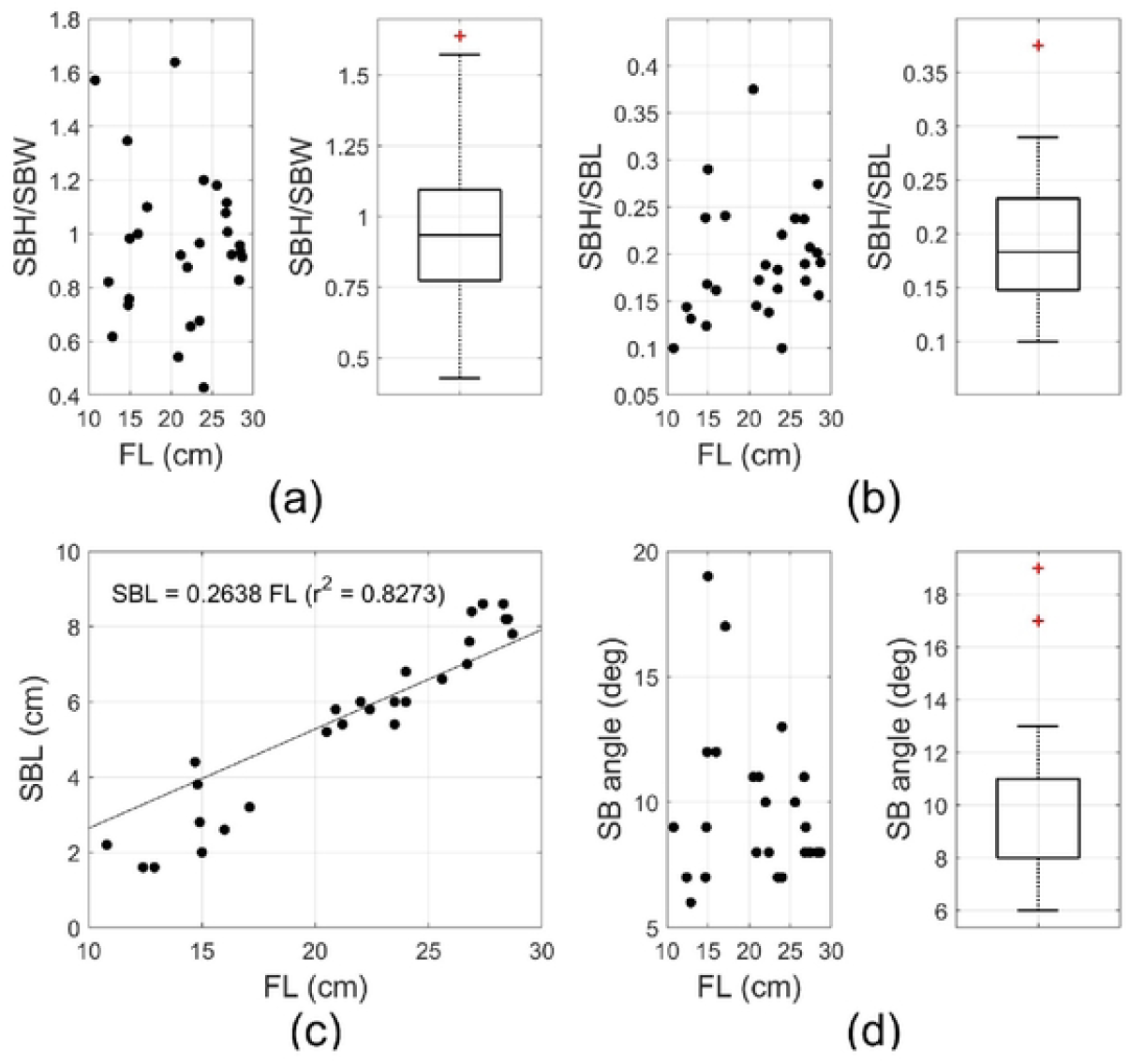
Length (SBL), height (SBH), width (SBW), and angle (SB) of swimbladder of chub mackerel specimens. (a), left: ratio of swimbladder height and width against fork length (FL); right: boxplot of SBH/SBW. (b), left: ratio of swimbladder height and fork length; right: boxplot of SBH/SBL. (c) swimbladder length and fork length, showing regression line with 95% confidence interval. (d), left: swimbladder angle vs fork length; right: boxplot of swimbladder angle.

## 4. Discussion

### Chub mackerel TS sample specimens

Given the species’ characteristics, chub mackerel TS experiments to date have mostly been conducted using dead specimens due to the difficulty of working with live individuals [19,20]. We were able to conduct TS experiments on various size groups of living chub mackerel by making use of active set-net fishing in Tongyeong, allowing for the capture of live specimens and a short transport time to the experimental facility. When multiple specimens are placed in a cage for experimentation, there can be an overlap in group sizes, which could potentially lower the reliability of mean size estimation and the TS relationship. However, since chub mackerel are a schooling species, they might exhibit abnormal swimming behaviors when isolated in a tank. We therefore placed up to five specimens at a time in a cage to acquire acoustic signals in a stable swimming state. To ensure no overlap in lengths within size groups, specimens were visually selected in the acclimation tank, minimizing group size deviation to 1.1 cm or less and resulting in almost no overlap between groups (one-way ANOVA, F = 244.0, p<0.05). Cage method experiments on European whitefish (*Coregonus lavaretus*) have been conducted for five size groups over a length range of 5–59 cm, with a tolerance for a mean size range within groups of 15 cm, showing significant variation [13]. This indicates that a wider size range within a group leads to a relatively wider TS range. In contrast, our study, although covering a smaller range of 10.8–28.3 cm, obtained TS signals for nearly identical size groups within an average size variation of only 2 cm, yielding more precise TS range estimates.

Among methods for fish TS experimentation, the caged method allows obtaining the TS of fish swimming relatively naturally compared to the tethered method, where fish are fixed with fishing lines and artificially subjected to changes in swimming posture angles [15]. In the present study, we simultaneously acquired TS signals and free-swimming posture angles, resulting in single modes and significant between-group differences for most groups, indicating that the data were obtained in a stable state.

### Chub mackerel swimbladder characteristics

In this study, the morphological parameters of fork length and swimbladder length, width, and height in chub mackerel showed a high correlation. Fish swimbladders are morphologically categorized into five types [29]. The SBH/SBL ratio, SBL/FL ratio, and swimbladder tilt angle (mean ± SD) of chub mackerel used in the experiment were 0.191 ± 0.060, 0.245 ± 0.055, and 9.6 ± 3.0°, respectively, suggesting a classification as round swimbladders (with the standard given as 0.2, 0.282, and 6°, respectively). Chub mackerel swimbladders were previously reported as showing a long extension, bending below the vertebral column, with a volume-to-body ratio of 3.2%, length ratio of 25%, and posture angle of 11.9° [20]. The reported range of body length of 18 specimens was 12.98∼22.17 cm, with a mean of 16.08 ± 3.15 cm (mean ± SD); the SBL range was 1.45∼6.63 cm, with a mean of 3.64 ± 1.49 cm; and the swimbladder tilt angle was 1.7° to 14.0°, with a mean of 8.26° ± 3.62° [7]. Although both of these studies lack SBH/SBL results, the SBL/SL findings of our study also classify our specimens as having round swimbladders.

### Chub mackerel swimming posture angles

In this study, the mean swimming posture angles for groups of chub mackerel ranged widely from −10.4–9.6°. In comparison, chub mackerel observed in a smaller rectangular tank (82 cm × 28 cm × 28 cm) displayed narrower variances in swimming posture angles of −4 ± 4° [30]. It seems that many fish with swimbladders have close to horizontal mean swimming angles, as has been found for cod (−4.48 ± 16.28° [31]), capelin (3.88 ± 18.48° [32]), caged saithe (−0.98 ± 48° [14]), hoki (11.88 ± 29.98° [33]), *in situ* Pacific saury (−1.1 ± 15.4°) and Japanese anchovy (−1.3 ± 20.8° [34]). The present study observed a range of swimming posture angles among chub mackerel that was broader than in other species. It was notable that angles changed sharply when fish ascended or descended within the cage. Previous reports suggest that even small juveniles might not show natural swimming behavior within a 3 m^3^ cage [35]. These results suggest that natural swimming posture angles that would be encountered in the field are difficult to represent in captivity, suggesting that further observations and comparative analysis using underwater cameras are required.

### Chub mackerel TS

The major factor in TS versus fish length regressions is tilt angle, acoustic frequency, fish length, and depth at the time of measurement [36–39], which adhere to the following order of influence: tilt angle > frequency > fish length > depth [40]. The caged measurement method can result in variance in swimming posture angles due to restricted movement within the narrow cage. In the present study, the group-specific TS values mostly showed a unimodal distribution, ranging widely from 10–30 dB. Additionally, changes in swimming posture angles became more pronounced as group size increased. TS measurements of European whitefish using the cage method also showed a wide distribution of 17–19 dB for larger sizes and 10–13 dB for smaller sizes, as well as a widening range of TS differences as group size increased [13]. Variations in TS using the tethered method and models showed significant differences even under the same alterations in swimming posture angles as size increased [41]. When comparing average group-specific TS values converted to b_20_ based on changes in swimming posture angles, the highest b_20_ values were observed at an average swimming angle of −4.1° across all frequencies, with b_20_ decreasing as the variation in swimming angles increased. While the tethered method allows for control over swimming posture angles while measuring TS, thereby providing clear TS values for each swimming angle, in this study, we compared mean swimming posture angles and b_20_ across groups. Therefore, the variation in the swimming angle must be examined to correctly interpret the TS data.

Previous studies on *S. japonicus* TS were all conducted using dead specimens (Table 2). Reported TS measurements at *b*_*20*_ using the tethered method included −64.1 dB at 25 kHz and −65.5 dB at 100 kHz for lengths of 23.0–26.8 cm [19] and −67.2 dB, −69.9 dB, −66.9 dB, and −71.1 dB for lengths of 26.2–38.3 cm at 50, 75, 120, and 200 kHz, respectively [20]. A previously reported KRM model used for frequencies of 38, 70, 120, and 200 kHz for *S. japonicus* TS (FL=15.4–26.2 cm) b_20_ yielded −66.02, −66.50, −66.00, and −67.35 dB, within 1 dB of the present study’s results [22], while another study reported values 5–6 dB lower, indicating significant outcome differences based on model parameters [7].

**Table 2.**
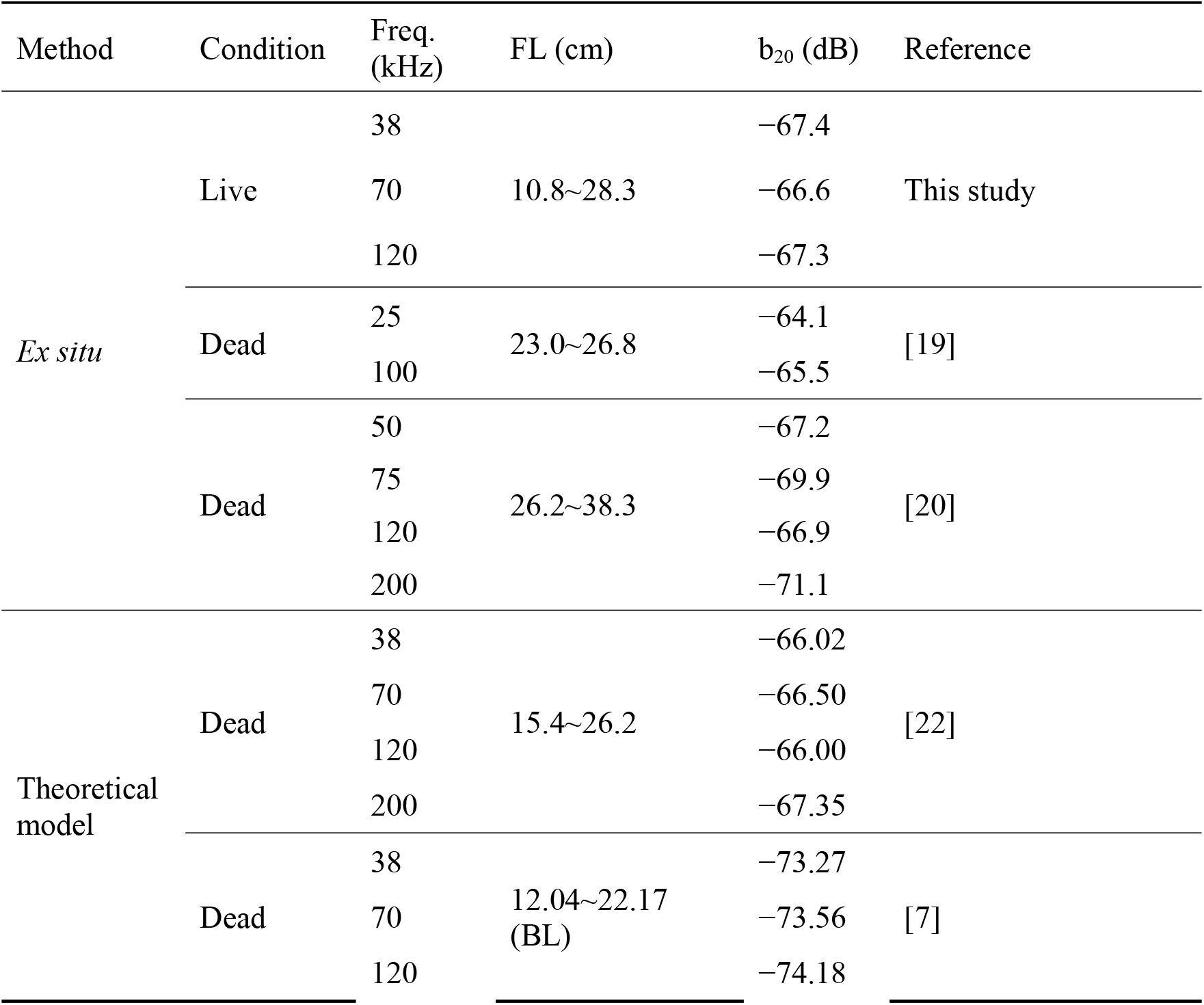
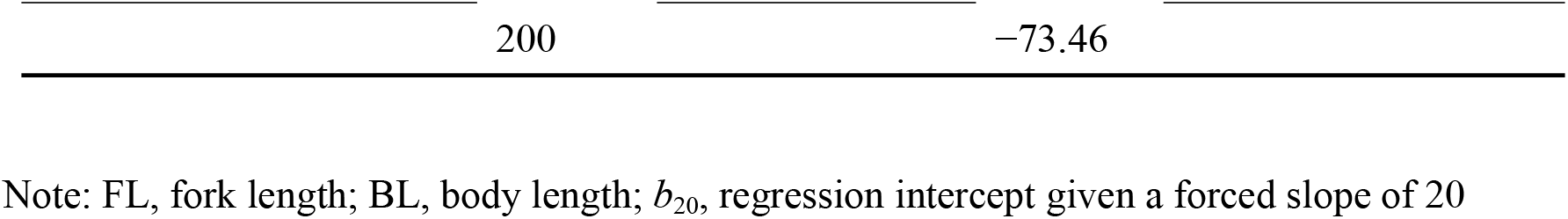
Comparisons of TS-FL relationship in chub mackerel by acoustic frequency and experimental method in the present and previous studies.

We discovered a high correlation between group-specific TS and length for chub mackerel across all three tested acoustic frequencies, with the optimal model slope consistently being close to 20. This indicates that the horizontal cross-sectional area of the swimbladder, as that in the generalized fish model, is proportional to the square of the length in this species, affecting TS values [15]. Additionally, the decrease in slope with increasing frequency suggests that the variation in TS due to swimming behavior becomes more pronounced at higher frequencies. In conclusion, we successfully derived a TS-L relationship for various sizes of living chub mackerel, and the findings will be utilized as foundational data for evaluating the density and stock of chub mackerel.

## Author Contribution

Conceptualization: Euna Yoon, Hyungbeen Lee; Data curation: Euna Yoon, Yong-Deuk Lee, Yong Jin Choo; Formal analysis: Euna Yoon. Funding acquisition: Jung Nyun Kim; Investigation: Euna Yoon, Hyungbeen Lee; Methodology: Euna Yoon, Hyungbeen Lee, Jeong-Hoon Lee; Writing– original draft: Euna Yoon; Writing– review & editing: Euna Yoon, Hyungbeen Lee

## Acknowledgements

We would like to thank the researchers of the Fisheries Resources Research Center for helping with the experiment. We are grateful to the two anonymous reviews that helped improve this paper.

## Ethics Statement

This study was carried out in strict accordance with the recommendations in the Guide for the Care and Use of the National Institutes of Fisheries Science. The protocol was approved by the Committee on the Ethics of the National Institutes of Fisheries Science (Protocol Number: 2022-NIFS-IACUC-24).

